# Anterior cingulate is a source of valence-specific information about value and uncertainty

**DOI:** 10.1101/087320

**Authors:** Ilya E. Monosov

## Abstract

Expectations of rewards and punishments can promote similar behavioral states, such as vigilance, as well as distinct behavioral states, such as approach or avoidance. However, the cortical circuits that underlie this behavioral diversity are poorly understood. In a Pavlovian procedure in which monkeys displayed a diverse repertoire of reward, punishment, and uncertainty related behaviors not mandated by the task, I show that many anterior-cingulate (ACC) neurons represent expected value and uncertainty in a valence-specific manner, for example about either rewards or punishments. This flexibility may facilitate the top-down control of many reward- and punishment-related actions and behavioral states.

## Introduction

Actions to maximize rewards and minimize threats or punishments are thought to be controlled by a region of the prefrontal cortex (PFC) called the anterior cingulate cortex (ACC). But how this control is accomplished is unclear. Generally, there are several views of how ACC controls motivated behavior (1–9). The first posits that the ACC neurons rank-order outcomes (such as rewards and punishments) by their utility or value. Then, the resulting “signed” value signal from ACC is read-out by other brain areas to guide action towards the best possible option or outcome (2, 5–8, 10). This theory envisions that many single ACC neurons ought to carry information about both rewards and punishments, and rank-order predictions of these outcomes according to their value (for example, as has been observed in the lateral habenula and amygdala (11–13)). A second theory posits that ACC controls negative-affect, threat, and pain driven behaviors. This theory suggests that many ACC neurons ought to display a preference for punishment-related information (14–17). A third theory suggests that the ACC is primarily engaged in deploying attention to monitor task-performance and detect errors in our predictions and actions (18–25). This theory predicts that many ACC neurons ought to be activated by salient, engaging, and cognitively demanding situations, irrespective of their valence. Finally, a fourth theory, based on recent imaging studies, identifies ACC as a central hub for processing information about uncertainty and other exploration- and learning-related variables, such as the value of foraging (26–30). Several views suggest that ACC could support some or even many of these functions (1, 3, 4, 6, 7, 25, 30–39). However, to date, it remains unclear how the ACC would do so.

Among the several difficulties in reconciling these views, one is that it is not known whether the same ACC neurons can signal the value and uncertainty of predictions about rewards and punishments. To address this issue, here single neuron activity was recorded from ACC while monkeys experienced certain and uncertain predictions about rewards and punishments cued by well-learned single visual fractals that served as conditioned stimuli. This approach has been used by previous studies to isolate value and uncertainty signals in the activity of single neurons from choice and learning-related modulation of subjective value, and timing and outcome uncertainty (12, 13, 40–43).

The results of the experiments suggest that the ACC contains distinct groups of neurons that process appetitive, aversive, salient, and uncertain information, as well as other neurons that multiplex information about certain and uncertain predictions of different valences. The data also suggest that many ACC neurons are sensitive to context (36, 44), resulting in encoding of motivationally relevant information on short- and long- time scales (34, 45). Overall, the data caution against a unified view of ACC and suggest that the ACC may contain circuits that contribute to the top-down control of a wide-range of behaviors and actions in distinct but complementary ways.

## Results

To test how single ACC neurons signal information about reward, punishment, and uncertainty, two monkeys were conditioned with an appetitive-aversive behavioral procedure that contained two separate contexts, or *blocks*. One block contained 12 trials in which three visual fractal objects (conditioned stimuli, CS) predicted rewards (juice) with 100, 50, and 0 *%* chance. The second block contained 12 trials in which three visual fractal CSs predicted punishments (air puffs) with 100, 50, and 0 % chance (Figure 1A). The monkeys did not have to fixate the CSs to complete the trial (Methods). The design was such that, 100% reward CS theoretically had the highest value in the reward block, and 0% punishment CS had the highest value in the aversive-block. Hypothetical encoding strategies of reward and punishment predictions are shown in Figure 1B (value exclusively for punishment- or reward-associated CSs, green and blue line, but not both; general value for both punishment- and reward-associated CSs, purple line; predicted intensity of outcome regardless of whether it’s a reward or punishment, red line).

**Figure 1.**
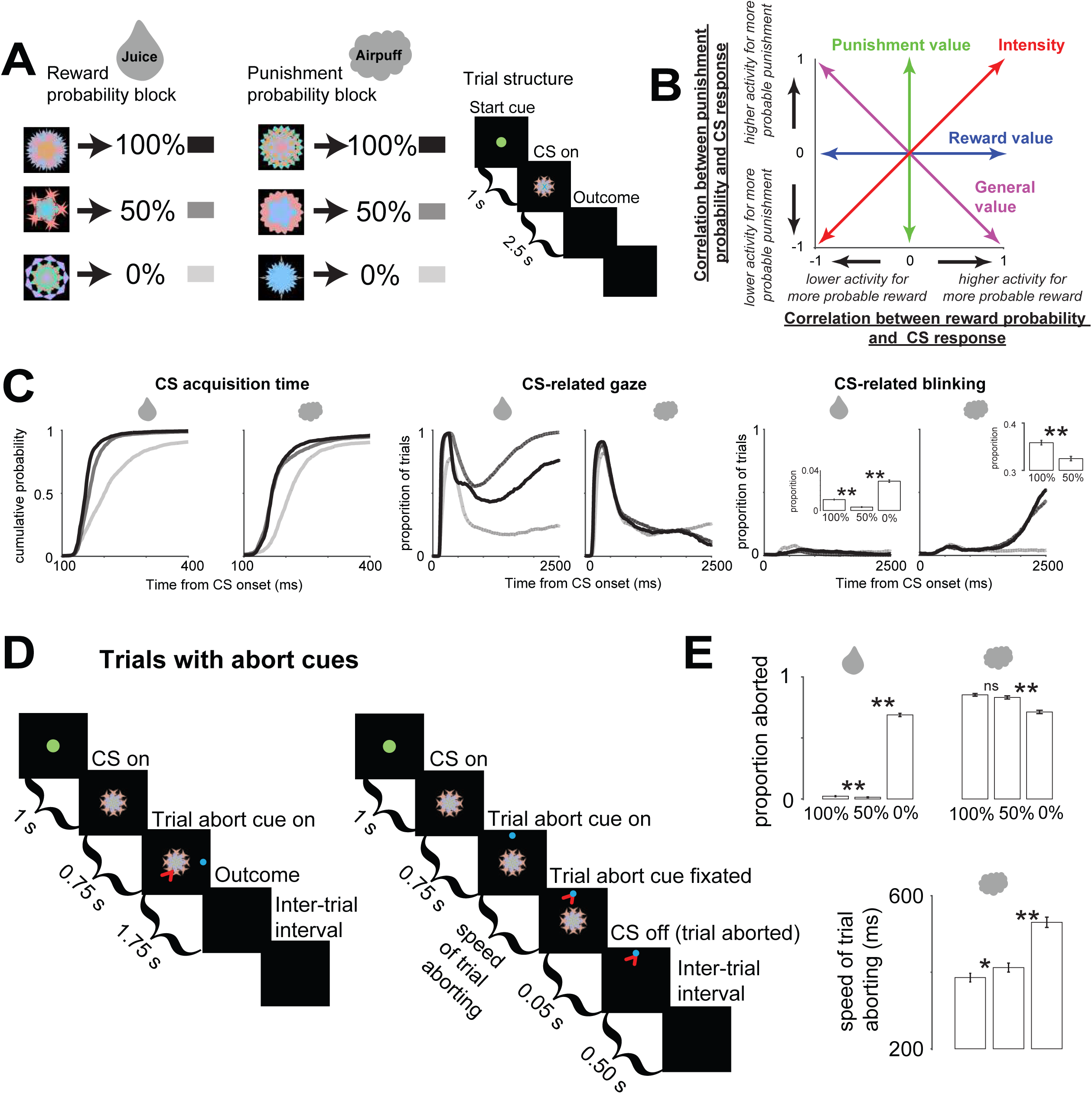
The reward-punishment behavioral procedure. (**A**) Monkeys experienced two types of distinct blocks of trials in which three visual fractal conditioned stimuli (CSs) predicted rewards and punishments with 100, 50, and 0% chance. The reward block consisted of 12 trials in which reward was possible, and the punishment block consisted of 12 trials in which punishment was possible. (**A-right**) Structure of a single trial. CS could appear in the center (as shown) or peripherally, 10 degrees to the left or right of center. (**B**) Theoretical valence coding strategies in the reward-punishment procedure. (**C**) Monkeys’ behaviors were motivated by reward, punishment, and uncertainty. (**Cleft**) Cumulative probability functions of peripheral-CS gaze acquisition time. In reward or punishment block, the speed of CS acquisition was correlated with outcome probability (Spearman’s rank correlations; p<0.01). (**C-middle**) Proportion of trials the monkeys oriented to the location of peripheral CSs shown across the entire CS epoch for reward- and punishment-block trials. During the last 1000 milliseconds of the trial the monkeys’ gaze was preferentially attracted to the 50% reward CS (p<0.01; rank sum test). (**C-right**) Proportion of trials the monkeys blinked shown across the entire CS epoch for reward- and punishment-block trials. Insets show mean proportion of time monkeys blinked during the last 500 ms of the CS epoch. (**D**) In distinct blocks we included an abort cue during approximately one half of the trials (Methods). Structure of trials with an abort cue. (**D-left**) – trials in which the monkeys did not abort (eye position is schematized by the red arrow); (**D-right**) – trials in which the monkeys aborted the trial. (**E**) Proportion of trials aborted in the reward block (top-left) and punishment block (top-right). The speed of trial aborting decreased with increasing punishment probability (bottom). The black-gray color legend for (**C and E**) is in (**A**). Error bars indicate standard error. Single asterisks indicate significance at a 0.05 threshold and double asterisks indicate significance at 0.01 threshold.

As in our previous studies (46, 47), after > 1 month of training, the monkeys understood the behavioral procedure. Their CS-related anticipatory mouth movements (e.g., licking of the juice spout) and anticipatory blinking were correlated to the probability of reward and punishment (Spearman’s rank correlations; p<0.01), respectively. While these anticipatory responses were related to the expected value of the CSs, other behaviors, such as gaze, were driven by the absolute expected value of the CSs (often called motivational intensity or salience (48)) and by outcome uncertainty. Particularly, the monkeys’ gaze was initially drawn to the CSs associated with the most probable outcome (100 % CSs; Figure 1C – left), irrespective of valence. Target acquisition times during 100% CS trials were faster when compared with all the 50% and 0% trials, within either reward or punishment block (p<0.05; rank sum tests). Within the reward block, target acquisition times were faster during 100% reward CS trials than during 50% reward CS trials (p<0.05; rank sum tests). However, later in the trial the monkeys’ gaze was most strongly attracted towards the 50% reward CS (Figure 1C – middle; rank sum test; p<0.01). We could not reliably observe CS-related gaze behavior during the last second of the punishment-predicting CSs because the gaze signal was quenched by defensive blinking (Figure 1C – right).

To study punishment, it is crucial to verify that the outcome or unconditioned stimulus is aversive. It could be that blinking behaviors in Figure 1C reflect conditioning, but not aversion. To address this issue, we utilized distinct reward and punishment blocks that contained abort cues during about one half of the trials (Methods). This paradigm is a type of active avoidance that is often used to test the aversiveness of cues, outcomes, and contexts in humans and experimental animals (49–53). If the monkeys made a saccade to the abort cue, the trial was aborted (Figure 1D). Because these reward- and punishment- blocks alternate, the optimal reward-driven strategy would be to rapidly abort every trial in which the 100% and 50% reward CSs were not presented. In contrast, the monkeys’ aborting behaviors were influenced by reward, punishment, and uncertainty. In the punishment block, monkeys actively aborted punishment-predicting CSs more often than the 0% punishment CS, confirming that the air puff was an aversive outcome (Figure 1E). While initially 100% punishment CS was associated with faster target acquisition than 0% CS (suggesting that it was more motivationally salient because it strongly attracted gaze; see ref 48), later in the trial (when abort cues were presented), monkeys aborted the 100% CS faster than 50% and 0% punishment CSs. The data show that the monkeys’ motivation to abort was positively related to the probability of punishment.

Also, monkeys aborted 50% reward CSs less than 100% reward CS trials (though, note that the proportion of aborted trials during either 100 or 50% reward CSs was extremely low). This decrease in abort-error rate is consistent with the observations in Figure 1C (middle) that the 50% reward CS captured attention (despite having a lower expected value than 100% reward CS), reducing the number of saccades to the location of the abort-cue.

In sum, the behavioral data suggest that monkeys utilized different representations (or encoding strategies) of rewards and punishments to influence their behaviors. Next, we asked if the ACC contains distinct representations of rewards and punishments or if ACC neurons encode rewards and punishments with a common currency, such as a general value signal (Figure 1B). The locations of ACC neuronal recordings are shown in Supplemental Figure 1 and match the locations of neuronal recordings in previous studies of macaque ACC (5, 9, 35, 54, 55).

The neuronal recordings revealed (*1*) that many ACC neurons represent expected value in a valence-specific manner, displaying greatest sensitivity to the probability of either rewards or punishments; and (*2*) also that many ACC neurons prefer uncertain CSs in a valence-specific manner, often displaying preference for either reward or punishment uncertainty.

First, to summarize the valence encoding strategies of ACC neurons, we performed correlation analyses for each recorded neuron (n=329) which assessed the relationship of CS-responses and outcome probability (Methods; Supplemental Figure 2–3). The correlation coefficients are shown in Figure 2A-inset. Most neurons that displayed significant correlations with outcome probability (Spearman’s rank correlation; p<0.05) did so in a valence specific manner, for either reward or punishment probability.

**Figure 2.**
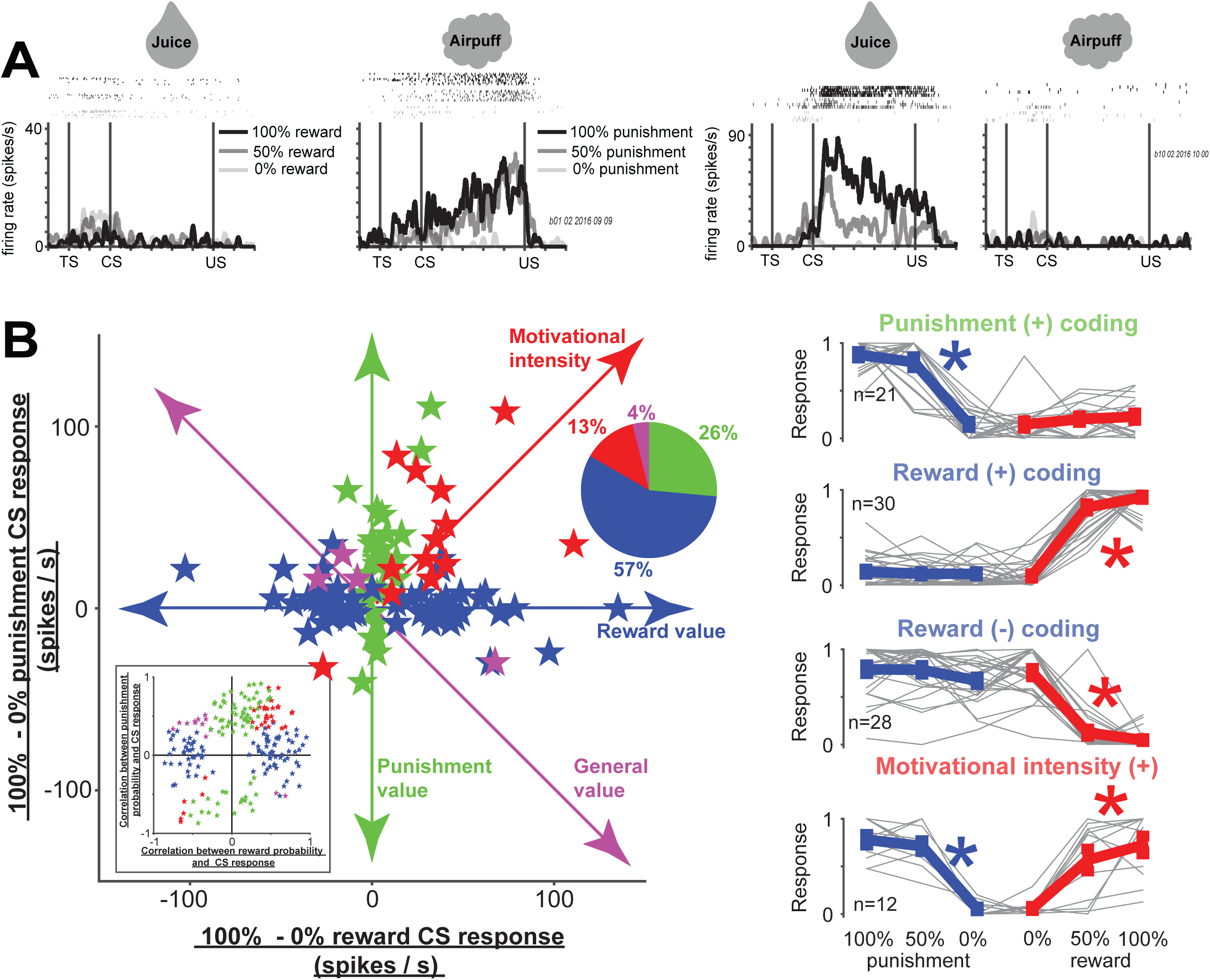
Many anterior cingulate (ACC) neurons signal outcome probability in a valence-specific manner. (**A**) Two single ACC neurons, one that signals punishment probability (left) and one that signals reward probability (right). (**B**) Scatter plot shows the difference in CS responses for 100% and 0% CSs in the reward block (x-axis) versus difference in CS responses for 100% and 0% CSs in the punishment-block (y-axis). Green – neurons displayed significant differences between 100% and 0% CSs in the punishment block only, blue - neurons that displayed significant differences between 100% and 0% CSs in the reward block only, red – neurons that displayed significant differences between 100% and 0% CSs in both blocks and both differences were either positive or negative, and purple - neurons that displayed significant differences between 100% and 0% CSs in both blocks and the differences were of different sign. Colors in pie chart (inset) correspond to colors in scatter plot and in Figure 1B. Uncertainty-selective neurons are studied separately in Figure 3. (**B-inset**) Correlation coefficients of all recorded neurons that displayed significant correlations with either reward or punishment probabilities, or both. Significance of each correlation was tested by 10,000 permutations (p<0.05). (**B-right**) Single neuron responses (gray) shown separately for 4 major groups of outcome probability coding neurons in the scatter plot in (**B**). Single neurons’ CS responses were normalized to the maximum CS response; from 0 to 1. Red asterisks indicate that the neurons’ responses varied significantly within the reward block (Kruskal-Wallis test; p<0.01) and displayed significant correlation with outcome probability (Spearman’s rank correlation; p<0.01); blue asterisks indicate that the neurons’ responses varied significantly within the punishment block (Kruskal-Wallis test; p<0.01) and displayed significant correlation with outcome probability (Spearman’s rank correlation; p<0.01).

Two valence-specific example neurons are shown in Figure 2A. The first neuron displayed greatest excitation to punishment-predicting CSs and did not differentiate amongst 100, 50, and 0% reward CSs. The second neuron displayed greatest excitation to reward-predicting CSs (strongest to 100% reward) and did not differentiate amongst 100, 50, and 0% punishment CSs.

Population analyses (Supplemental Figure 3) and a qualitative examination of single neurons’ responses (Figure 2A) suggested that many ACC neurons signal value in a valence-specific manner. To assess this on a population level in ACC, we analyzed the differences in neuronal responses for good outcomes and bad outcomes in the reward (x-axis in Figure 2B) and punishment (y-axis in Figure 2B) blocks. These analyses confirmed that most value coding neurons display valence-specific responses (blue and green colored data in Figure 2). The second most common type of ACC neurons signaled the motivational intensity or unsigned value of the CSs (red in Figure 2). Few neurons showed a generalized value signal that rank-ordered the reward or punishment predictions according to their expected value (100>50>0% rewards and 0>50>100% punishments; purple dots in Figure 2). The time courses of neuronal responses among these neuronal types are shown in Supplemental Figures 5–6.

Many neurons’ prediction-error outcome responses also tended to be valence specific. For example, many ACC neurons distinctly signaled reward omissions, reward deliveries, punishment omissions, and punishment deliveries (Supplemental Figure 3). Interestingly, the value of the CSs and their associated outcomes were often signaled by different neurons (Supplemental Figure 3). To summarize, the data thus far suggest that ACC can encode the values of predictions and deliveries of rewards and punishments in distinct population of neurons.

ACC neurons discriminate among tasks (36), behavioral contexts (44), and integrate task-related information over long time scales (34, 45). Hence, it was important to assess if ACC contains valence specific neurons in a behavioral procedure in which reward and punishment CSs are presented in the same context or block of time. The results of this control experiment verify that many ACC neurons co-vary with the probability of only reward or only of punishment, when their predictions are experienced within a single block of trials (Supplemental Figure 7). However, this result does not indicate that ACC valence specific neurons are context insensitive or that they are not sensitive to the statistics of salient events over long time scales. In contrast, the data in the two block appetitive/aversive procedure indicate that monkeys and ACC neurons were highly sensitive to the nearing of their preferred context over many trials (Figure 3). Here, behavioral and neuronal responses to the trial start cue were analyzed. Though trial start cues were presented 3.5 seconds before the trial’s outcome would be experienced by the monkeys, their anticipatory orienting behavior (the duration it took them to foveate the trial start cue) was significantly faster in the reward versus punishment block (Figure 4 – left). Also, during the punishment block the speed of target acquisition changed as a function of trial number, slowly decreasing as the reward block neared (p<0.001; rho=0.12). Similarly, reward and punishment sensitive neurons signaled the nearing of their preferred blocks. Reward-sensitive neurons displayed gradual and systematic changes in the aversive block in anticipation of the reward block (Figure 4 – middle). And, punishment-sensitive neurons displayed systematic changes in the aversive block in anticipation of the aversive block (Figure 4 – right). Hence, ACC neurons can signal valence specifically or non-specifically (Figure 2) and can signal the nearing of their preferred contexts or events over long time scales (Figure 3).

**Figure 3.**
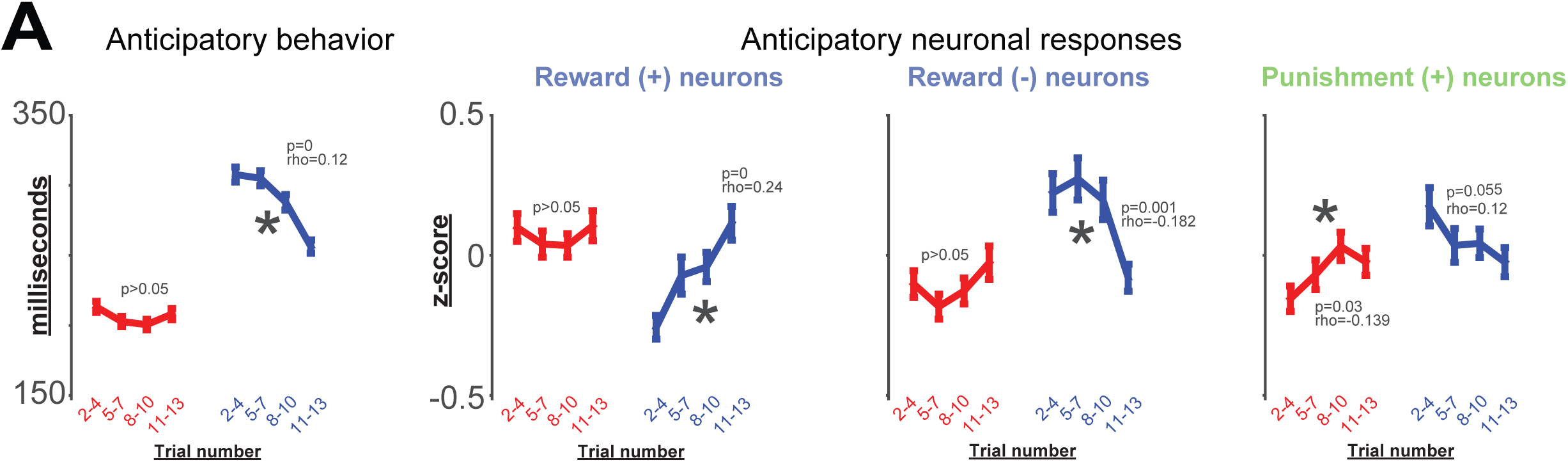
Monkeys and ACC neurons anticipate the nearing of their preferred block. (**A**) Reward and punishment blocks of 12 trials were separated into four sub-blocks. The blocks were separated relative to the monkeys’ knowledge about which block they were in. For example, the reward block starts after the first reward CS trial after which the monkeys knew they were in the reward block. It ends during the first punishment CS trial (because the monkeys had not yet seen the punishment CS). Hence, the block trial numbers in this figure are numbered 2 through 13. The average times of when monkeys foveated the trial start cue (relative to the time of the trial start cue presentation) are shown for the four sub-blocks in the reward block (red, left) and aversive block (blue, right). The results of a Spearman’s rank correlations assessing the relationship of subblock number and speed of orienting to the trial start cue are indicated above each block. Asterisk highlights when a correlation was significant. (**B**) Responses of the three groups of valence-specific neurons identified in Figure 2 during the last 500 milliseconds of the trial start cue epoch. Conventions are the same as in (**A**). In all 4 plots, the data in the reward block (red) differed from the data in the punishment block (blue; rank sum test; p<0.05).

**Figure 4.**
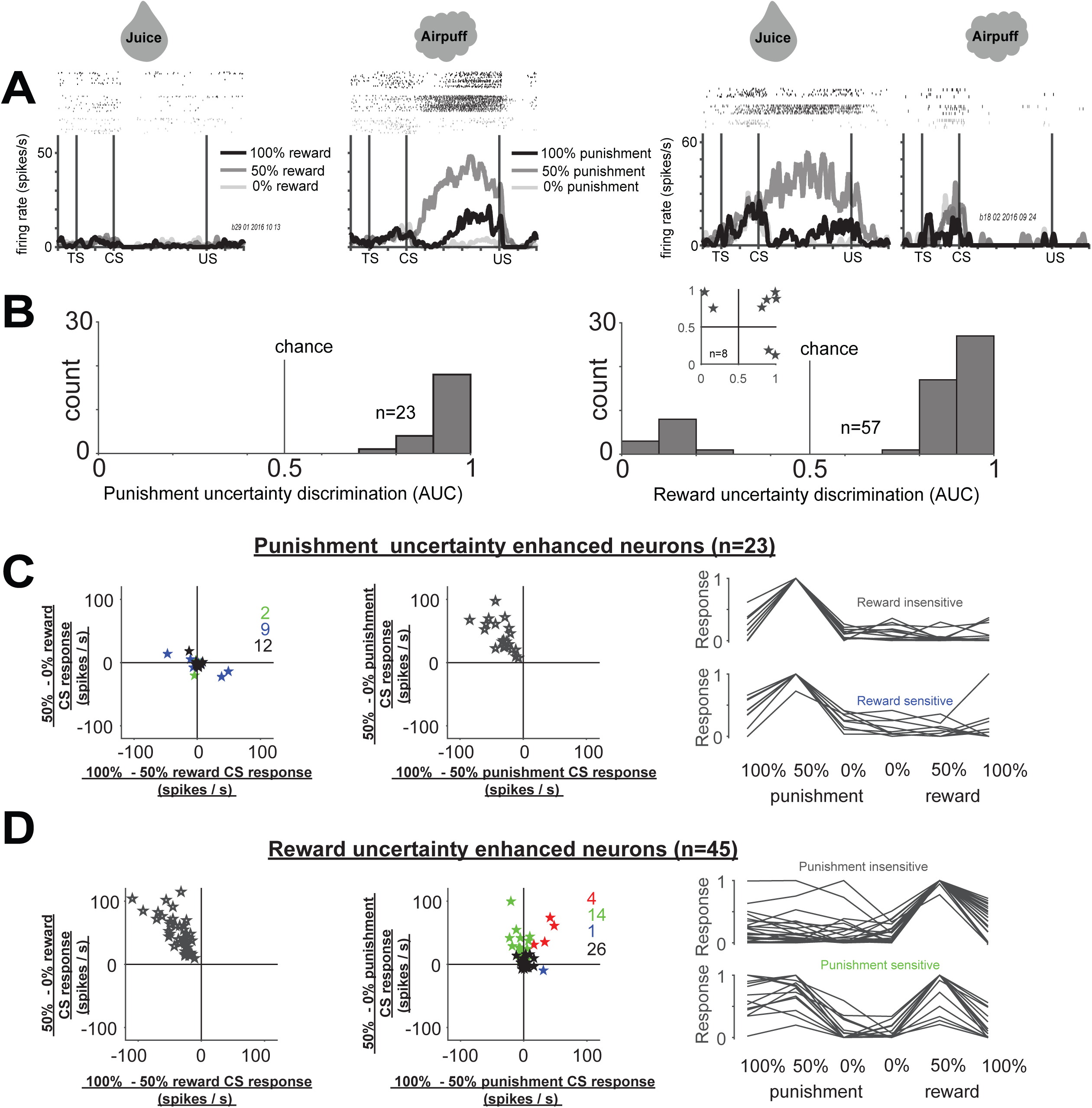
Distinct coding of reward and punishment uncertainty by many single ACC neurons. (**A**) Two single ACC neurons, one that signals punishment uncertainty (left) and one that signals reward uncertainty (right). (**B**) Single neuron discrimination indices of punishment and reward uncertainty selective neurons (top and bottom). Inset shows discrimination of 8 neurons that displayed uncertainty selectivity in both reward (x-axis) and punishment blocks (y-axis). Discrimination indices were obtained by ROC analyses (Methods). Values below 0.5 indicate selective suppression, indices above 0.5 indicate selective enhancement. (**C**) Punishment-uncertainty enhanced neurons in the reward block (left-scatter plot) and punishment block (right-scatter plot). Scatter plots show, for individual neurons, the differences between 50% versus 100% CS responses (x-axis) and 50% versus 0% CS responses (y-axis). Red stars, Neurons showing significant differences in CS response in both 50–100% comparison and 50–0% comparison. Green stars, Neurons showing significant differences in only 50–0% comparison. Blue stars, Neurons showing significant differences in only 100–50% comparison. Black stars, neurons showing no significant differences. Single neuron and average CS responses (right) for two major groups of punishment uncertainty enhanced neurons (black and blue circles on the left). (**D**) Reward-uncertainty enhanced neurons. Conventions are the same as in (**C**).

Next, we assess if and how single ACC neurons encode uncertainty. In contrast with the hypothesis that ACC encodes economic or general “common currency” values, some recent neuroimaging studies have highlighted the ACC as a central hub for processing outcome uncertainty and guiding behaviors aimed to reduce this uncertainty. However, single neuron evidence for uncertainty processing in ACC has been missing.

We found 88 uncertainty selective neurons in the ACC (Methods), some of which responded selectively for punishment uncertainty. An example neuron is shown in Figure 4A (left). This neuron did not respond to reward predicting CSs or reward outcomes. In the punishment block, the neuron was most strongly activated by 50% punishment CS and this signal persisted until the trial outcome. Amongst uncertainty selective neurons, 23/88 neurons were selectively excited by punishment uncertainty but not reward uncertainty (Figure 4B; Supplemental Figure 8). 0 were selectively inhibited by punishment uncertainty (Figure 4D). Among the punishment-uncertainty neurons, 11 neurons displayed significant variability among the reward block CSs (Figure 4D – left; blue dots) but there was no obvious trend for positive or negative reward value coding amongst them. These results show that some ACC neurons signal punishment uncertainty in a selective manner, and that a minority of them can also multiplex this uncertainty signal with information about rewards.

Consistent with the observations that ACC can signal task-related information in a valence specific manner (Figure 2), we also found neurons that preferred reward uncertainty but not punishment uncertainty (Figure 4B). An example neuron is shown in Figure 4A (right). This neuron responded to the 50% reward CS and maintained tonic firing until the time of the trial outcome (reward or no reward). It was not modulated by punishment predicting CSs. 45/88 uncertainty selective neurons were excited by reward uncertainty while 12/88 uncertainty selective neurons were inhibited. A 26/45 of reward uncertainty excited neurons also carried information about punishment probability; Figure 4B and Supplemental Figure 8). Among these, almost all were enhanced by increased probability of punishment (Figures 3C and 3E). A minority of uncertainty selective neurons (8/88) were selective for both reward and punishment uncertainty (Figure 4B – inset).

Like all other populations of uncertainty or risk coding neurons, ACC uncertainty neurons discriminated between 100% and 0% CSs in their preferred block (40, 46, 47, 56, 57). Within the aversive block, 9/23 punishment uncertainty neurons responded more to 100% than 0%, and 3/23 responded more to 0% than 100%. 22/45 uncertainty enhanced neurons responded more to 100% than 0% reward CSs, and 0/45 responded more to 0% than 100% reward CSs.

Outcome deliveries following uncertainty elicit prediction errors (48, 58, 59). However, the majority of ACC uncertainty neurons did not signal prediction errors (Supplemental Figure 9; and see Supplemental Figure 8 for qualitative observation). This observation provides further evidence for the notion that within ACC, there may be distinct groups of neurons tracking predictions and outcome- or feedback-related prediction errors (Supplemental Figure 3).

In sum, our findings show that ACC can signal uncertainty about either rewards or punishments in a valence-specific manner or to multiplex information about uncertainty and value in multiple manners (Supplemental Figure 10). Also, prediction errors following uncertain epochs were often signaled by distinct populations of neurons. Hence, the ACC may be a crucial hub for the flexible processing and integration of uncertainty- and value- related information.

Uncertainty can arise due to variability in outcomes or due to the possibility of making an error. For example, in our behavioral procedure aborting a reward associated CS is an error because it reduces the probability of reward on that trial to 0%. Though we observed few such errors (Figure 1), the anticipation of the abort cue resulted in increases of overt attention towards the reward-possible CSs and the activity of uncertainty neurons in ACC, but not value coding ACC neurons (Supplemental Figure 11). Therefore, ACC uncertainty neurons are sensitive to uncertainty arising due to internal processing (that increases attention (30)) as well as due to uncertain stimulus-outcome associations.

Several recent studies show that several subcortical brain regions in the septum and the striatum contain populations of neurons that signal the graded level of reward uncertainty (46, 47). Because these brain regions receive inputs from the ACC (60, 61), an important question then is, do ACC reward uncertainty neurons also signal graded levels of uncertainty and reward size? To answer this question, ACC reward enhanced uncertainty neurons were recorded while monkeys participated in the reward probability / reward amount behavioral procedures used in our previous studies (46, 47, 56, 62). The reward-probability block contained five objects associated with five probabilistic reward predictions (0, 25, 50, 75 and 100% of 0.25 ml of juice). The reward-amount block contained five objects associated with 100% reward predictions of varying reward amounts (0.25, 0.1875, 0.125, 0.065 and 0 ml). The expected values of the five CSs in the probability block matched the expected value of the five CSs in the amount block. The block design was used to remove the confounds introduced by risk-seeking related changes in subjective value processing of the CSs (40, 46, 47, 56)

Consistent with our previous reports using the same procedure, after conditioning, monkeys choices rank ordered the CSs in either block according to their expected values (Figure 5A) (46, 47, 56, 62) indicating they understood the meanings of the CSs.

**Figure 5.**
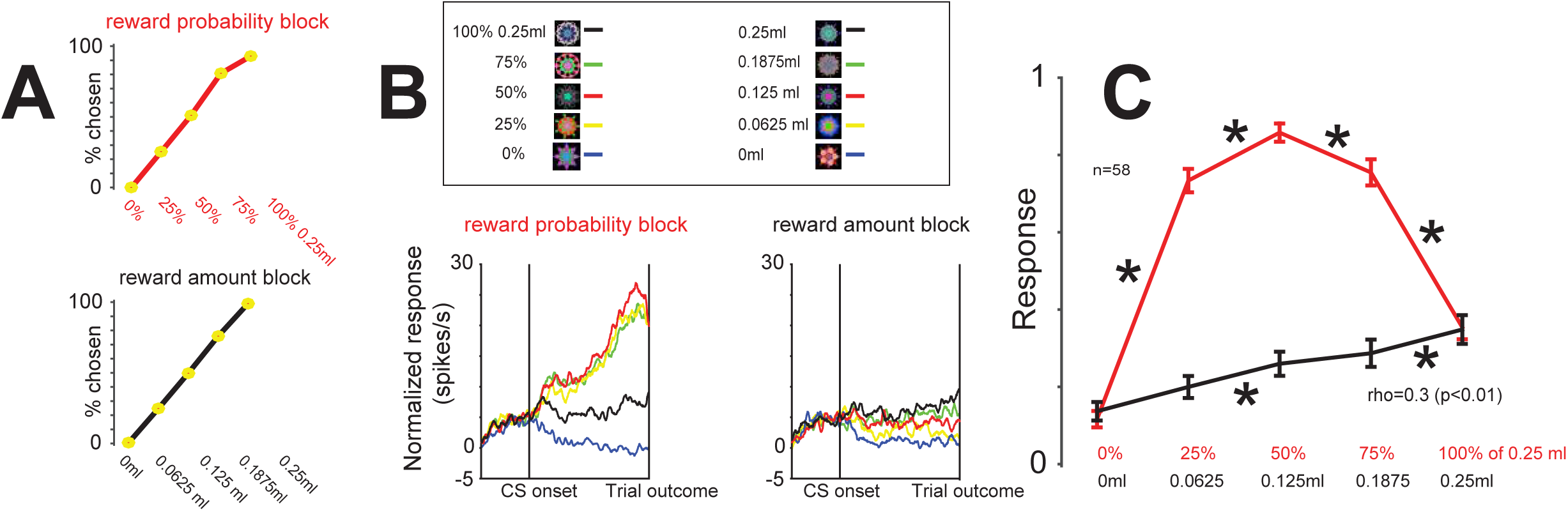
Reward uncertainty enhanced ACC neurons are sensitive to the level of reward uncertainty. (**A**) Monkeys’ choice preference between CSs associated with different reward amounts and probabilities. Choice percentage of a single reward probability CS (x-axis) versus all other reward probability CSs (top). Choice percentage of a single reward amount CS versus all other reward amount CSs (bottom). Data are compiled from a data set of 13407 choice trials. (**B**) Average responses of 58 reward uncertainty enhanced ACC neurons in the reward-probability block (left) and reward amount block (right). (**C**) Average normalized responses of the same neurons in the probability (red) and amount (black) CSs. Asterisks indicate differences between adjacent CSs (p<0.05; paired sign-rank test). The result of a Spearman’s rank correlation assessing the relationship of neuronal firing and reward amount in the reward amount block is indicated below. Before averaging, single neuronal responses were normalized to the maximum CS response, from 0 to 1, across the 10 different CSs. Error bars indicate standard error.

After training, 58 ACC reward uncertainty enhanced neurons were identified and studied in the single CS reward probability / reward amount behavioral procedure.

The online identification of reward uncertainty neurons was the same as in our previous studies (46, 47, 62).The average response of ACC reward uncertainty enhanced neurons is shown in Figures 5B-C. During the reward probability block the neurons responded most strongly to the most uncertain CS (50% CS), more weakly to 25 and 75% CSs. The same neurons displayed the weakest response to the certain CSs (0 and 100%). Consistent with observations in the reward / punishment behavioral procedure, these neurons also discriminated 100% versus 0% reward CSs (sign rank test; p<0.05). In the reward amount block, their responses encoded reward size in a roughly linear manner, displaying highest firing for the greatest CS associated with the greatest reward sizes (Figure 5C; rho=0.3; p<0.01; Spearman’s rank correlation). These data indicate that, on average, ACC reward-uncertainty neurons signal the level of reward uncertainty and can also signal reward amounts by shifting their tonic firing (Figure 5B).

## Discussion

To obtain rewards and avoid punishments we must detect them, estimate their uncertainty, and track and predict their value. Here, the data show that the anterior cingulate (ACC) has the capacity to mediate these functions by specifically and distinctly signaling information about the value and uncertainty of rewards and punishments, and to multiplex them. The multiple valence coding strategies in ACC may support its roles in the control of value-based action (2, 3, 7, 8, 31, 54, 63) and of uncertainty-driven processes such as learning and monitoring, risk-seeking, and the identification of novel or uncertain environments (1, 4, 29, 30, 35, 64).

The results are mostly inconsistent with several unified theories of ACC function. First, there was no indication that the majority of ACC neurons signaled aversive or noxious predictions or outcomes. Second, our data are inconsistent with the notion that many single ACC neurons encode a general value or common currency signal that generalizes across appetitive and aversive contexts. Specifically, very few ACC neurons displayed positive relationships with reward probability and negative relationships with punishment probability (or vice versa). Also, as in previous studies, within reward loss or reward gain trials, distinct neurons signaled predictions of reward gains or losses (7, 31, 65, 66). Since rewards and punishments can promote similar behavioral states, such as vigilance, or distinct behavioral states such as approach and avoidance, the heterogeneous valence-specific value signals in ACC may facilitate a flexible capacity for control of a wide range of behavior (1–4, 25, 34, 37, 54, 67) in distinct tasks and contexts (36, 44).

It was recently reported that that when rewards and punishments are bundled into a single stimulus (or option) that monkeys can choose to approach or avoid, neurons in the pregenual cingulate cortex (an area that is anatomically distinct from but related to regions studied in this report) were either correlated with approach behavior or avoidance (68). Interestingly, the avoidance-related neurons were also more activated when the monkeys’ decision reaction times were slow. The authors suggested that this effect may be related to the subjective conflict between the reward and punishment within a particular bundle (68). Along similar lines, Ebitz and Platt found that in a saccade task, distractor-induced conflict enhanced the activity of many neurons located within the ACC (25). Here, it was found that the possibility of an abort cue presentation during high valued trials (that monkeys rarely aborted) enhanced the activity of ACC uncertainty-sensitive neurons (Supplemental Figure 11). In these trials, the abort cue was an aversive distractor to which saccades ought not to be made. Together, these observations may suggest that task-conflict, the increased possibility of action-performance errors, and the possibility of prediction errors (following uncertainty), modulate similar groups of ACC neurons.

This study is the first to demonstrate a selective punishment-uncertainty signal in the brain. Punishment uncertainty neurons were often found in more anterior and ventral regions of the bank of the ACC (Supplemental Figure 1) while reward uncertainty neurons were found in all areas, but were particularly common in the dorsal extent of the ACC. Since the dorsal and ventral regions of the dorsal ACC have distinct projection patterns to the striatum, amygdala, cortex, and brain stem (60, 69–72), one possibility is that reward and punishment uncertainty are mediated by distinct circuits involving somewhat different (but overlapping) areas of ACC. This conjecture is further supported by a series of experiments in which we identified neurons in two anatomically distinct networks that selectively signal information about reward uncertainty, but not punishment uncertainty: the septal-basal forebrain network and the striatal-pallidal network (46, 47, 56, 62). Both, of these networks receive inputs from, and send direct and indirect inputs to, the ACC, and neither contains neurons that signal punishment uncertainty. Previous work has identified that the ACC may be important for monitoring decision uncertainty (64) and for information seeking (4, 73). It will be important to assess how different groups of neurons identified in this study, and throughout the subcortical uncertainty-related network, contribute to those functions.

The finding of a selective punishment uncertainty signal in the brain will provide new opportunities to study internal states such as anxiety and depression, which are thought to be strongly driven by persistent uncertainty about bad outcomes (28, 74–77). However, to move forward it will be crucial to identify precisely which brain regions receive and process punishment uncertainty signals from ACC and assess how the ACC interacts with the two reward-uncertainty selective networks to control reward uncertainty related action and internal states.

Several recent neurophysiological studies strongly suggest that ACC neurons display selectivity for contexts or tasks (36, 44). We observed that the reward/punishment preference of ACC value sensitive neurons did not change across blocks that contained active behavior to approach or avoid rewards and punishments, and blocks in which monkeys could not actively abort trials (abort-cues blocks versus no-abort cue blocks; Supplemental Figure 11). Also, when ACC neurons were studied in a control procedure in which rewards and punishments were predicted in the same context, many of the recorded ACC neurons again signaled value in a valence specific manner.

However, it can be argued that the differential behavioral selection (or action repertoire) related to the expectation of rewards or punishment imparts differences in contexts when subjects expect rewards or punishments (78, 79), and that the valence-specific value coding observed in ACC does not indicate that ACC neurons do not encode information in a context-sensitive manner. In fact, valence sensitive neurons predicted and anticipated the nearing of their preferred context over long time-scales (Figure 3). Also, on average, the CS responses of many reward and punishment sensitive neurons did not consistently differ across blocks in which abort cues may be presented versus blocks in which no abort cues were presented, suggesting that ACC neurons may have the capacity to adjust their responses relative to the values of the possible predictions within a given context or block of trials.

The remarkable capacity for valence - and context - sensitivities (36, 44) may delineate the ACC from some subcortical neurons such as dopamine and lateral habenula neurons. These populations only weakly discriminate rewarding and aversive contexts prior to CS onset (80) and, unlike some ACC neurons, are highly sensitive to the predictions and deliveries of both rewards and punishments. In contrast, recent evidence indicates that anterior 14c within the macaque ventromedial prefrontal cortex (vmPFC) and the dorsal raphe nucleus serotonin neurons (like ACC) display a capacity for context and valence specificity (41, 81, 82) by tonically and phasically signaling the valence of blocks or contexts. One possibility that requires serious investigation is that serotonin could play a crucial regulatory role in valence and context specific behavioral selection through raphe-to-prefrontal projections to the ACC and vmPFC.

Like all other conditioning studies, this study does not aim to dissociate action value from valence. To detect valence specificity in the activity of a single neuron, an experiment must use outcomes of different valences (78, 79). Rewards and punishments often elicit diverse and distinct actions and emotional or internal states (78, 79). This makes the dissociation of a valence-specific value signal from an action-value signal more difficult. Though ACC neurons, like neurons in the orbitofrontal cortex (43), did not display trial-by-trial correlations with conditioned responses (Supplemental Figure 12), they may control action by increasing or decreasing motivation in a valence, context, and action specific manner. Also, because this study sought to ask if ACC contained uncertainty selective neurons, this experiment was designed to decrease uncertainty that is unrelated to the uncertain-outcome conditioned stimuli. Hence, the behavioral procedure did not include choice trials in which uncertainty or risk can come about due to many factors (28, 40, 83–85). It will be important to understand the time course and dynamics of reward and punishment specific signals in ACC during choice behavior, particularly in economic decision making tasks in which rewards and punishments are combined into single choice-options.

Identifying neuronal mechanisms that facilitate flexible control of action and learning toward rewards and away from punishments remains a central pursuit of systems neuroscience. Here, we demonstrate a remarkable diversity and valence-specificity of ACC neurons that may in part provide a neuronal substrate for such capacity. The data suggest that the ACC has multiple distinct and cooperative functions in behavioral control related to the expectation and receipt of reward, punishment, and their uncertainty.

## Supplemental Figures

**Supplemental Figure 1.**
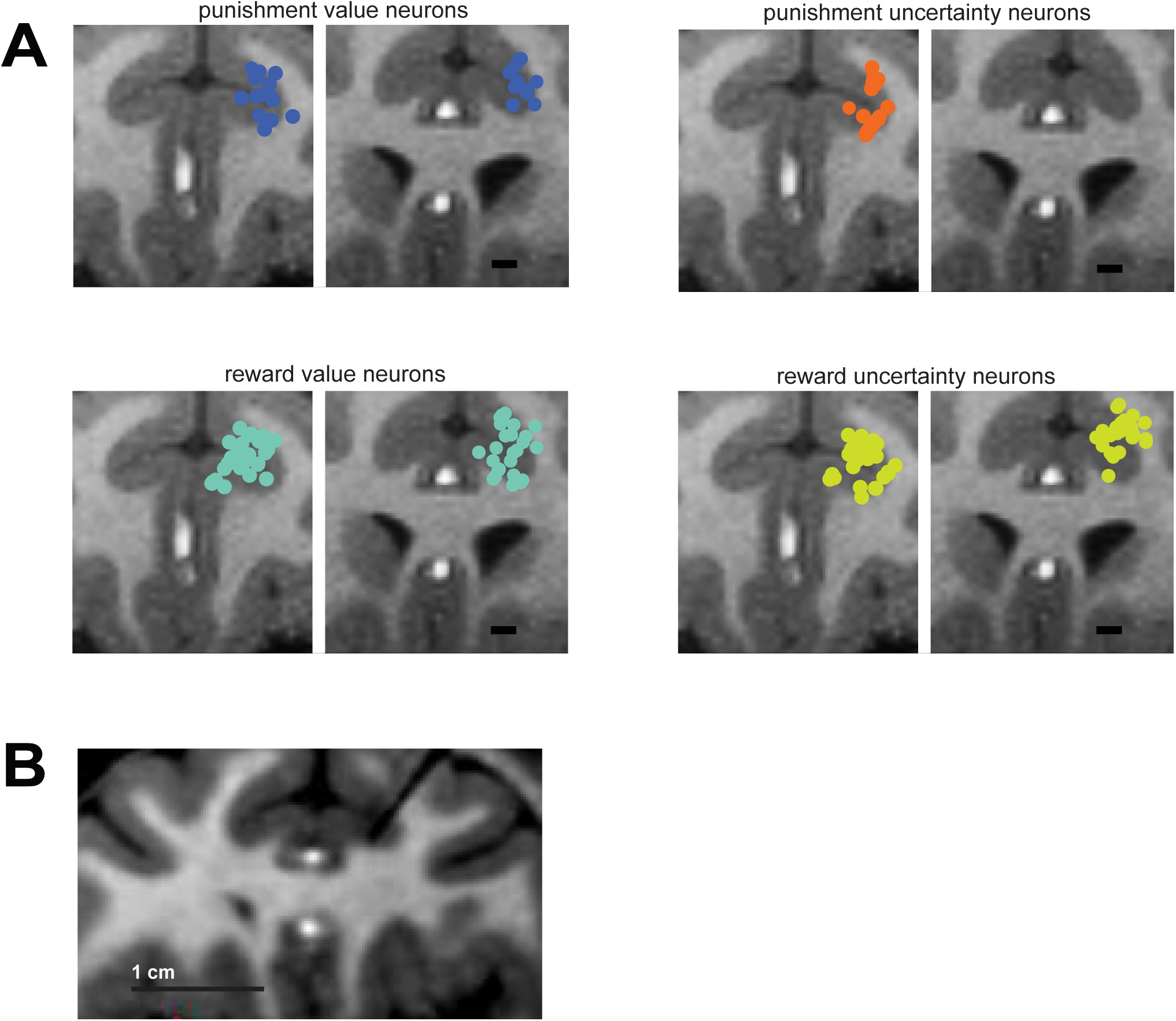
Estimated locations of reward value, punishment value, reward uncertainty, and punishment uncertainty ACC neurons. (**A**) The recording range was 6 to 17 mm from the center of the anterior commissure (AC), and also 3 neurons sampled 3 mm anterior to the AC were included in this study (which did not change any of the results). The anterior section on the left is 15 mm from the center of the AC and includes neurons recorded 17mm to 12 mm from the center of the AC. The posterior section on the right is 11 mm from the center of the AC, and includes the neurons sampled in the posterior extent of our recordings. It appears that different neuron types were mostly mixed within the ACC. One notable observation was the difference in the location of reward uncertainty and punishment uncertainty neurons. Punishment uncertainty neurons were found in the most anterior extent of our recordings and their locations were often within the ventral regions of the bank of the ACC, while reward uncertainty neurons were found widely throughout our recording area. (**B**) A coronal T1 magnetic resonance image confirming a recording location of a punishment uncertainty neuron in the ACC. The image was acquired with a tungsten electrode (FHC) at a recording location. The electrode’s shadow is the black line whose tip is in ACC.

**Supplemental Figure 2.**
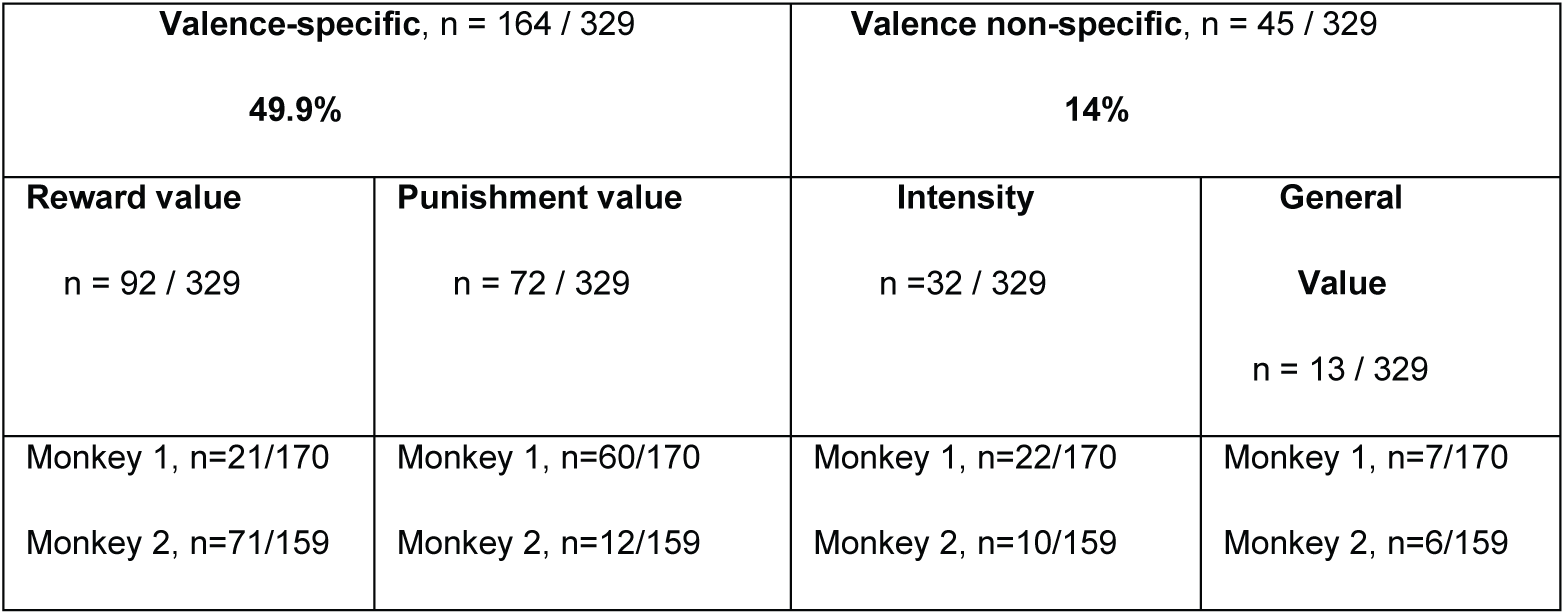
Summary of neurons’ correlations with predictions of reward and punishment. Data for single cells is shown in Figure 2-inset and Supplemental Figure 3A.

**Supplemental Figure 3.**
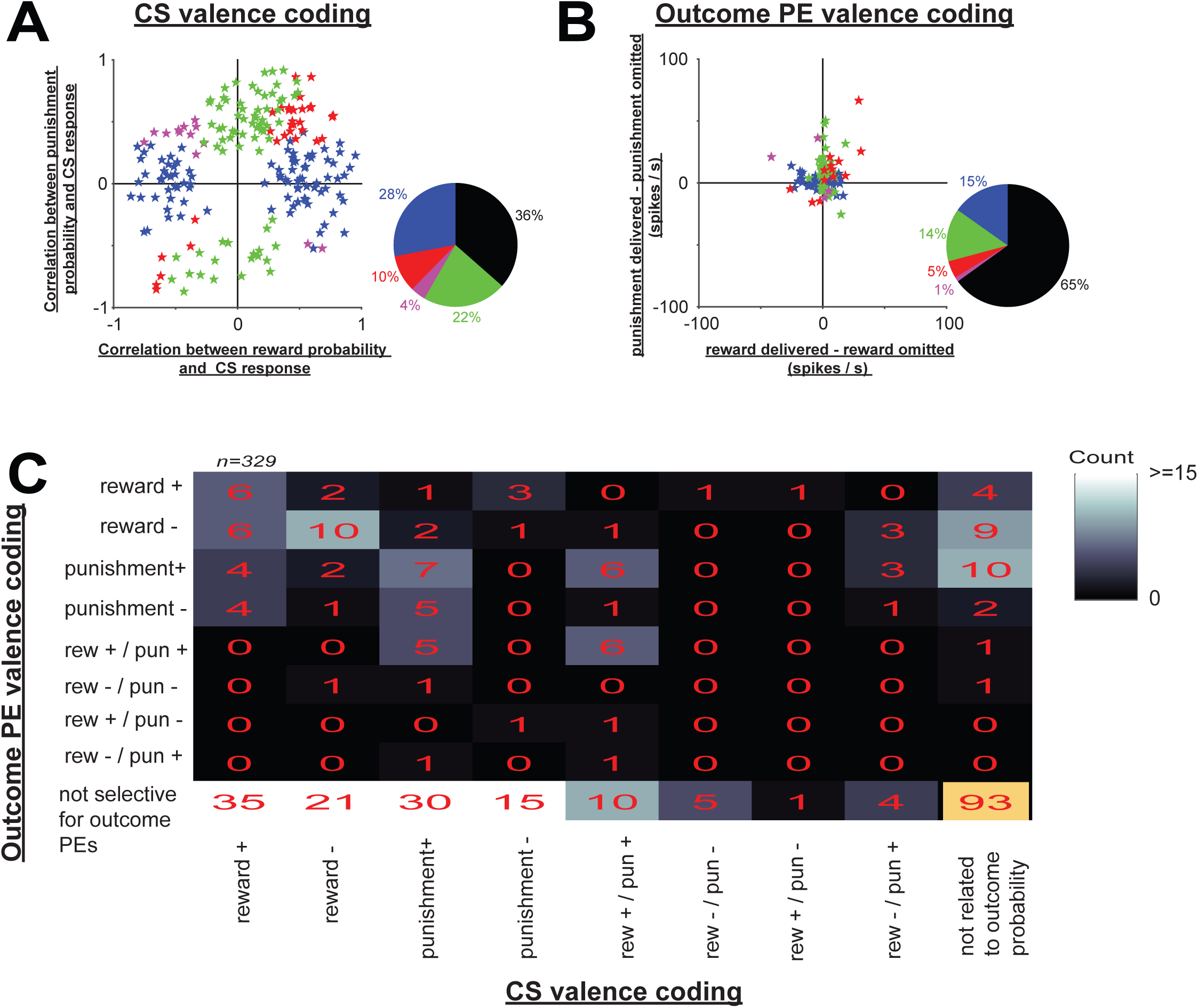
Coding of reward and punishments: CS responses and outcome prediction errors. (**A**) Correlation coefficients resulting in analyses for Table 1. Significance of each correlation was tested by 10,000 permutations (p<0.05). (**B**) Single neuron response differences between reward deliveries versus omissions following 50% reward CSs (x-axis) and differences between punishment deliveries versus omissions following 50% punishment CSs (y-axis). Significance was assessed by a Wilcoxon rank sum test (p<0.05). Colors in scatter plots and pie charts in **A-B** are the same as in Figure 2. (**C**) Counts of neurons displaying outcome PE coding (y-axis; determined from **B**) and CS sensitivity (x-axis; determined from **A**).

**Supplemental Figure 4.**
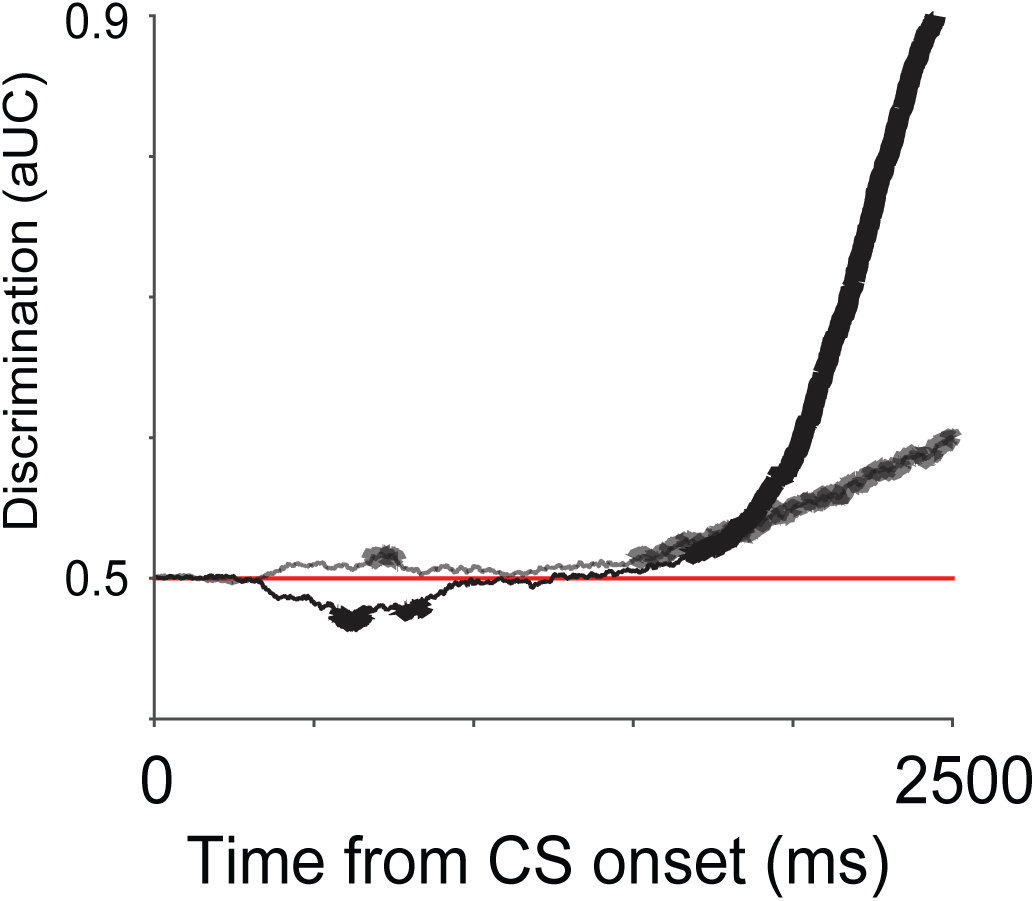
Expression of aversive conditioned responses in two monkeys. The results of an ROC analysis is shown in time comparing blinking during 100% and 0% CS punishment trials for Monkey 1 (black) and Monkey 2 (gray). Thicker lines indicate time points at which there was significant difference between 100% and 0% trials (rank sum tests; p<0.01 with Bonferroni correction for 2500 multiple comparisons for each millisecond in the CS epoch). Both monkeys’ blinking was strongly correlated with the probability of air puffs (Spearman’s rank correlations, p<0.01), but the difference between 100% and 0% CS trials was greater in Monkey 1. AUC – area under ROC curve.

**Supplemental Figure 5.**
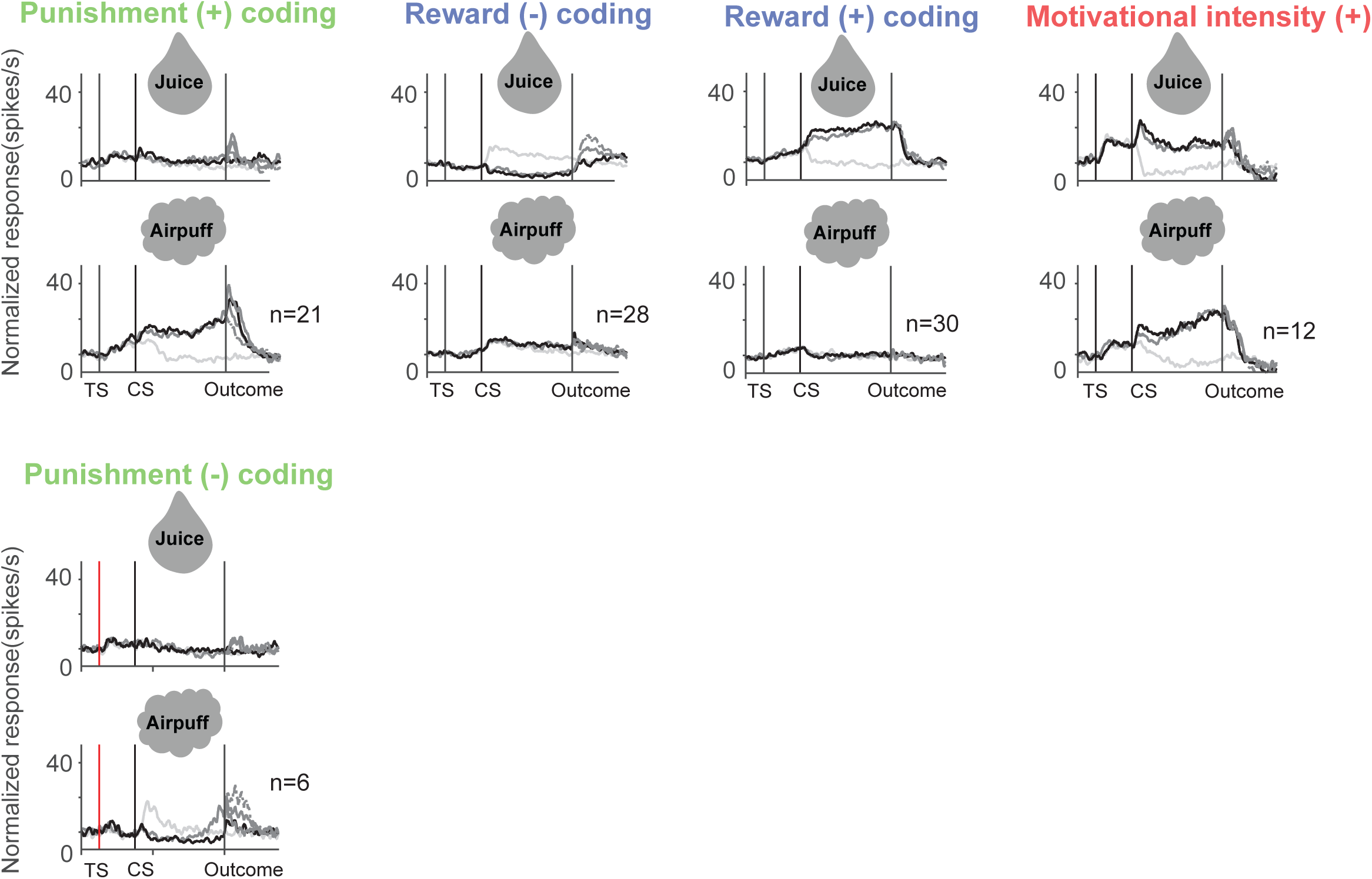
Neuronal activity of outcome probability value encoding neurons. Their activity in the reward (top) and punishment (bottom) blocks shown separately for 100, 50, and 0% CSs. Darkest traces show activity in 100% CS trials, lightest traces show activity in 0% CS trials (this is the same gray-black color scheme as in Figure 1). Dotted lines (after the outcome) represent outcome omissions.

**Supplemental Figure 6.**
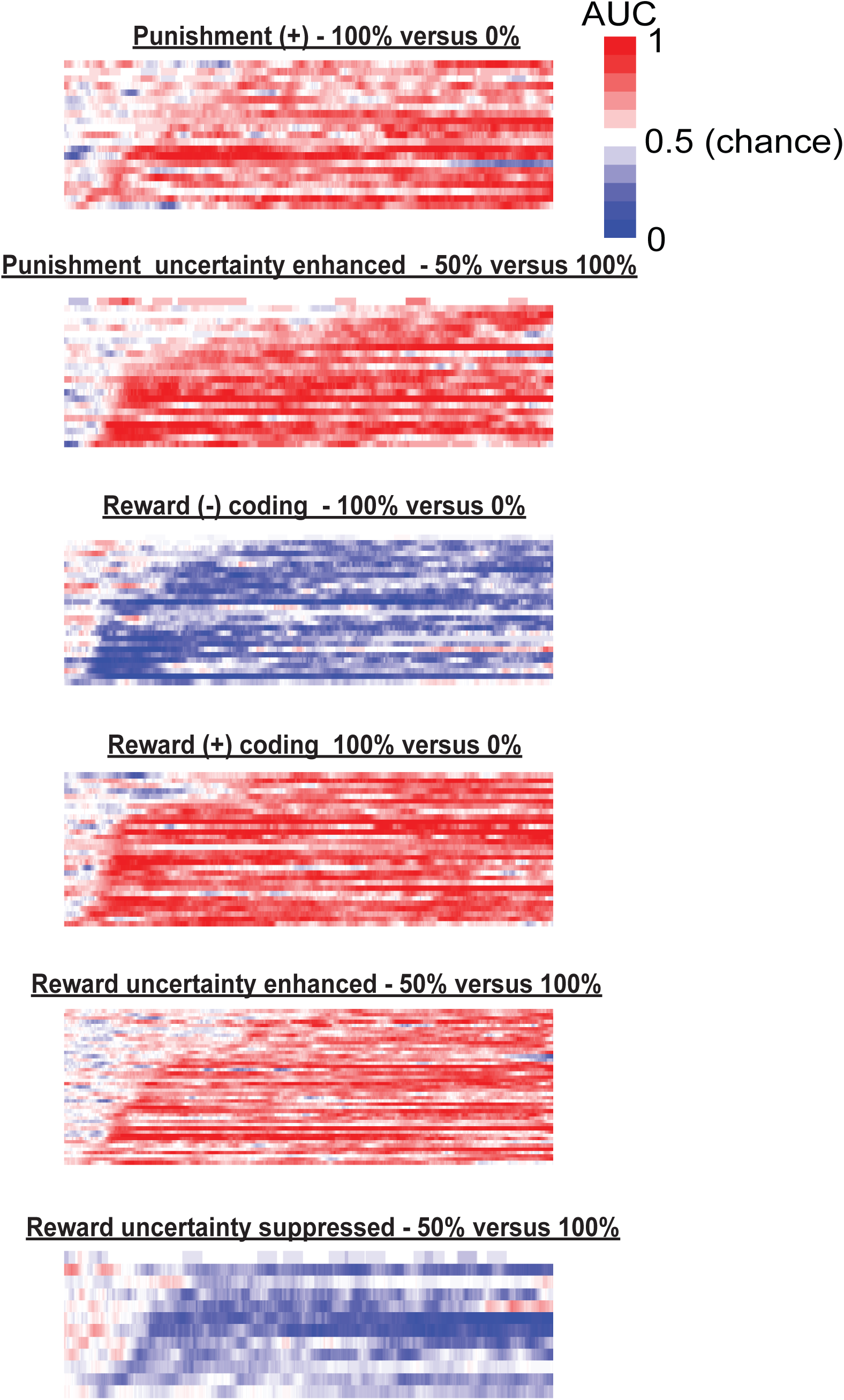
Time courses of value and uncertainty discrimination in single neurons. For each group of neurons (labeled above the heat maps), a receiver-operating characteristic (ROC) analysis was performed that compared spike density functions from CS onset to the time of outcome (2.5 s). Each line represents the results of this analysis for a single neuron during the entire CS epoch. As in our previous studies, for the value neurons the time course of value discrimination was assessed by an ROC analysis that compared 100% versus 0% CS trials in their preferred block. For uncertainty neurons, the time course of uncertainty discrimination was assessed by an ROC analyses that compared 50% versus 100% trials CS in their preferred block. ROC analyses were structured so that receiver-operating characteristic area values > 0.5 indicate that the activity in the 100% CS trials was greater than 0% CS trials for value neurons, and that 50% was greater than 100% CS trials for the uncertainty neurons; values < 0.5 indicate that the activity in the 100% CS trials was less than 0% CS trials for value neurons, and that 50% was less than 100% CS trials for the uncertainty neurons. AUC, area under receiver-operating characteristic curve.

**Supplemental Figure 7.**
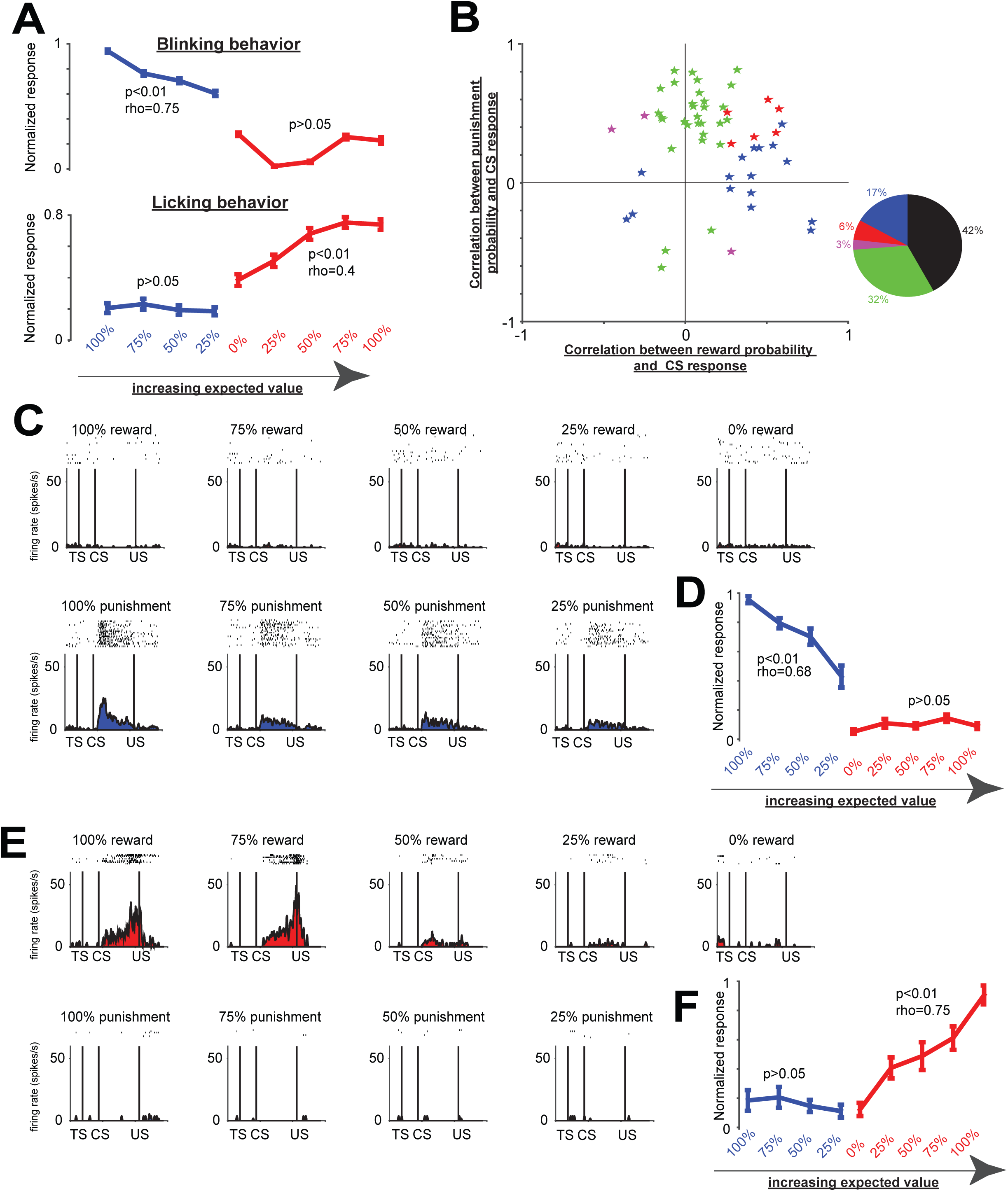
Single block appetitive / aversive procedure. (**A**) Conditioned responses to the presentation of 100, 75, 50, 25, 0% reward CSs and 100, 75, 50, 25% punishment CSs within a single block. Blinking (top) was correlated to the probability of punishment (rho=0.75; p<0.01), but not to the probability of reward (p>0.05). Though, as in Figure 1, blinking was least prevalent during reward uncertain conditions. Licking (bottom) was correlated to the probability of reward (rho=0.4; p<0.01), but not to the probability of punishment (p>0.05). (**B**) Correlations of single neurons with the probability of reward and punishment (same format as Supplemental Figure 3A). Pie chart shows the percentages of correlated neurons among all 95 recorded neurons. (**C**) The activity of a single punishment sensitive ACC neuron shown separately for all 9 CSs (formatting is the same as in Figure 2). (D) The average activity of 16 punishment enhanced neurons in ACC. Their correlation with reward and punishment probability is indicated above. (**E**) The activity of a single reward sensitive ACC neuron shown separately for all 9 CSs. (**D**) The average activity of 9 reward enhanced neurons in ACC. Their correlation with reward and punishment probability is indicated above. Error bars denote standard error. All analyses including the classification of neurons were the same as in Figure 2 (Methods).

**Supplemental Figure 8.**
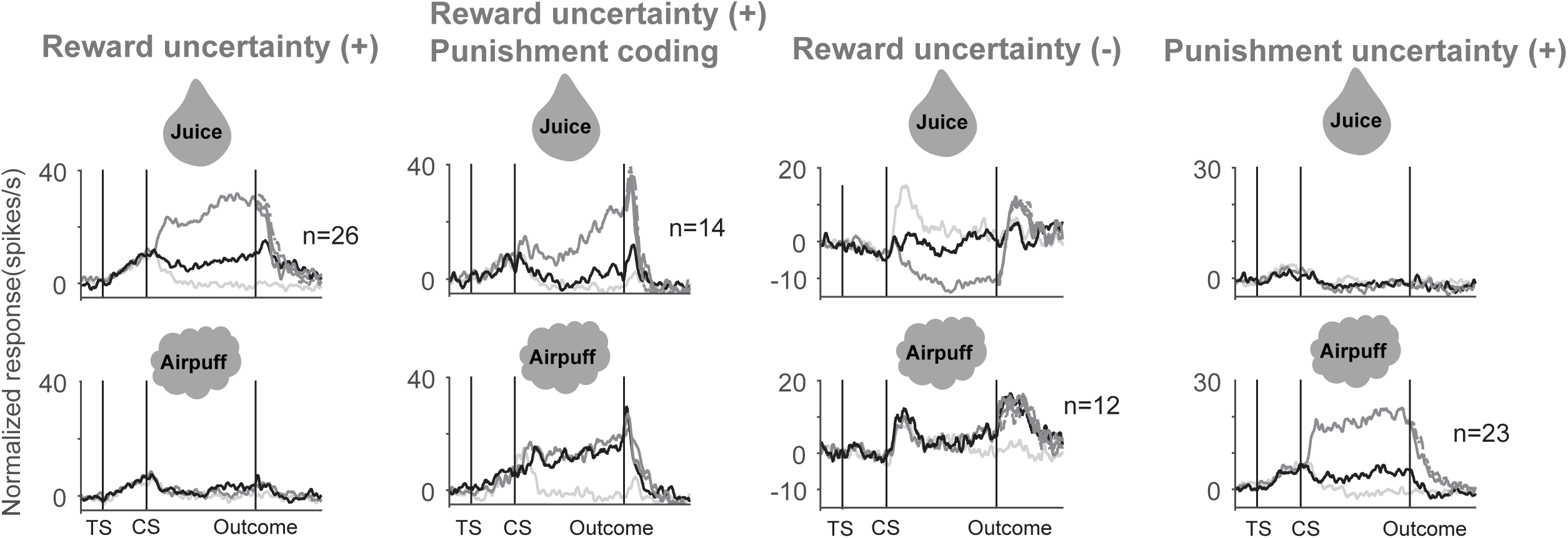
Neuronal activity of uncertainty selective neurons. Conventions are the same as in Supplemental Figure 3.

**Supplemental Figure 9.**
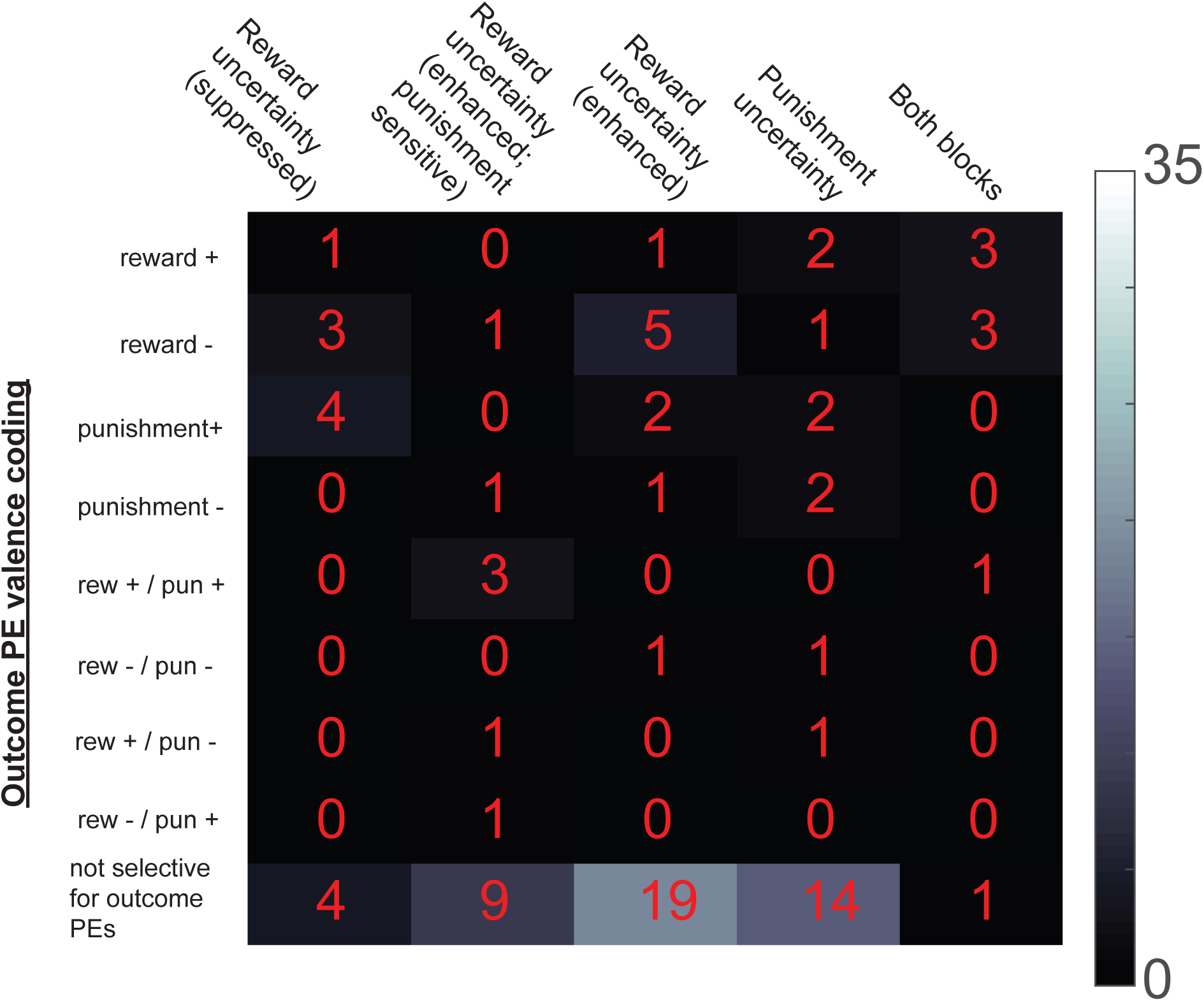
Neuronal counts of uncertainty neurons displaying outcome prediction error coding. Format is the same as in Supplemental Figure 3.

**Supplemental Figure 10.**
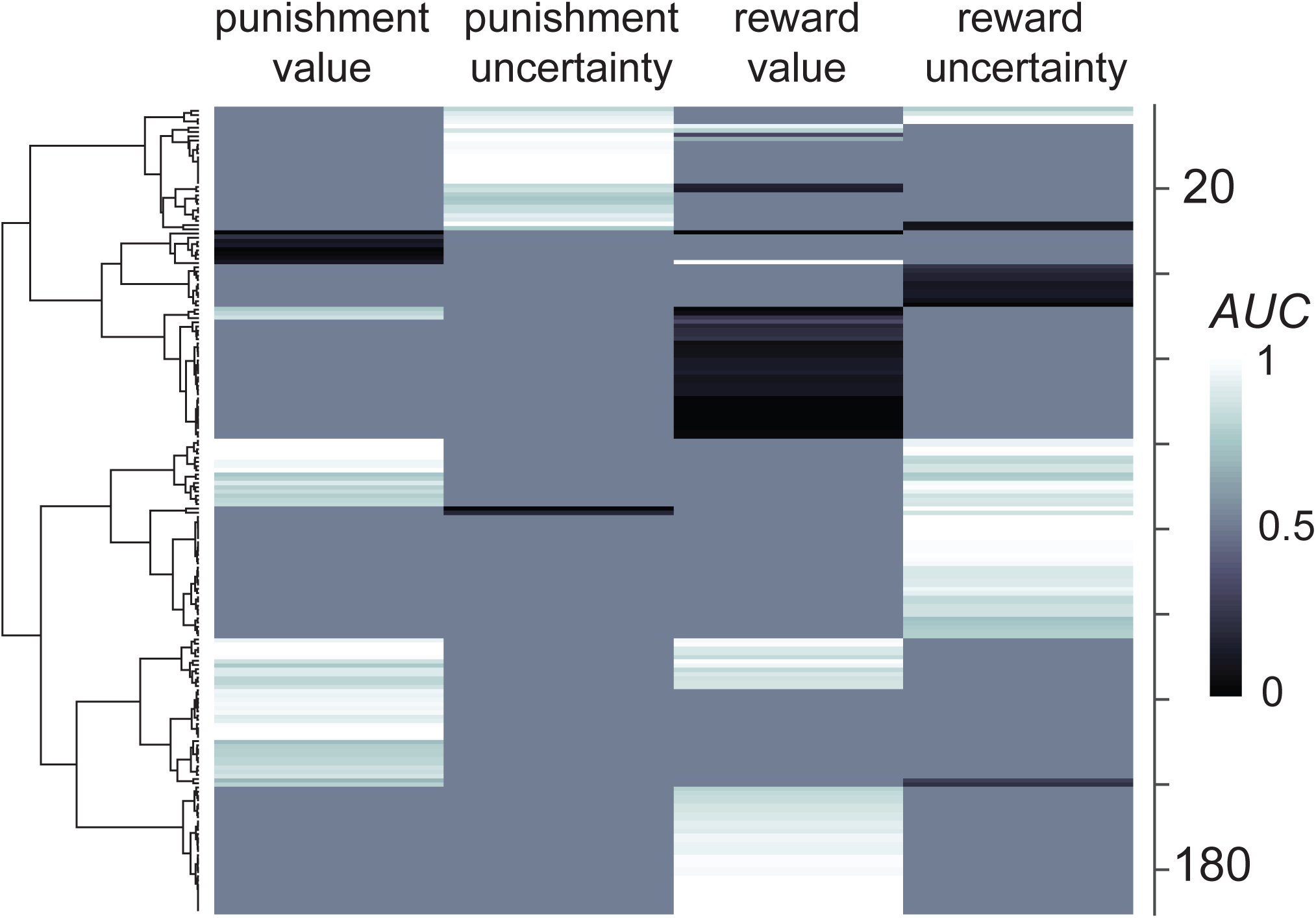
ACC neurons uncertainty and value coding. A visual summary of uncertainty and value sensitivity of all ACC neurons that displayed variance across the CSs (Kruskal Wallis test; p<0.01). Each neuron (shown as a row) contributes either an uncertainty or value discrimination index in each block. Matrix in this plot was organized by an unsupervised hierarchical clustering algorithm (Methods).

**Supplemental Figure 11.**
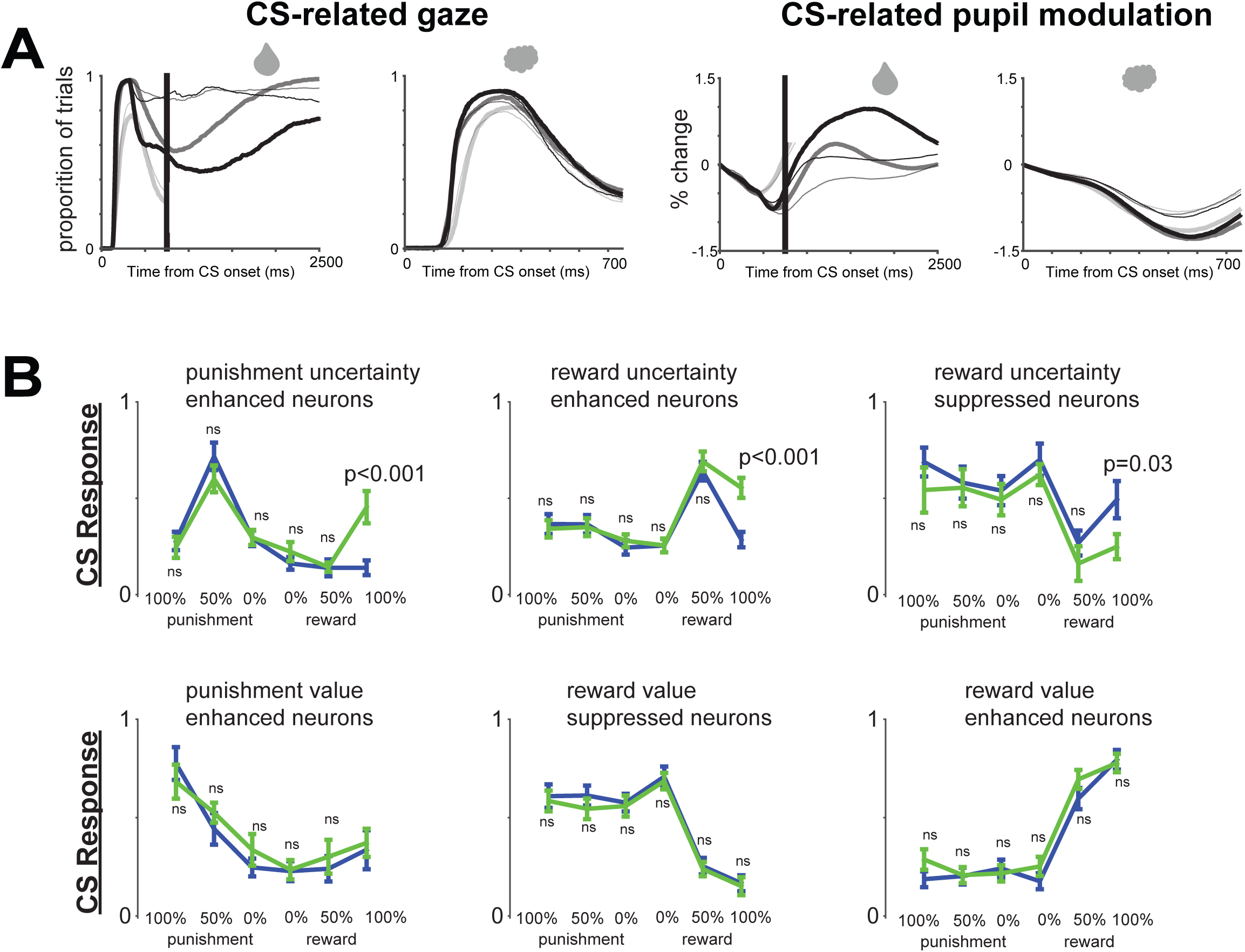
Behavioral and neuronal differences across blocks with and without abort cues. (**A**) Gaze behavior (left) and pupil dilation (right) is modulated by uncertainty and by abort possibility. Thick lines show behavior during blocks in which abort cues were never presented. Thin lines show behavior during blocks in which abort cues were presented 50 % of the time. Lines are colored as in Figure 1. Thick line at 750 ms following CS onset shows the time during which the abort cue would have been presented. Because the monkeys often aborted the non-rewarding trials and blinked during the later epochs of punishment-possible trials (Figure 1), we show behavior during 0% reward trials and 100, 50, and 0% punishment trials until the time of the abort cue would have been presented. After the time of the abort cue, we show behavior during the 100% and 50% reward trials in which monkeys did not abort. The possibility of abort cue presentation modulated gaze behavior (p<0.01; comparing reward-possible trials before the escape cues in abort-possible and no abort trials). In fact, monkeys could have looked away from the monitor to minimize the probability of error saccades after reward CSs were presented towards the abort cues. Instead, their overt attention increased towards reward CSs (100 and 50%). Pupil dilation (right) seemed to reflect gaze behavior (left), but with some lag. Both gaze and pupil dilation were affected by uncertainty (that was either due to the possibility of an abort cue during reward trials or by 50% reward predictions). (**B**) Neuronal activity before abort cues were presented (750 ms window following CS onset, before the abort cue) during blocks of trials in which abort cues were never presented (blue) versus blocks of trials in which the abort cue appeared during half of the trials (green) for 6 groups of ACC CS modulated neurons. Consistent differences were observed for uncertainty-sensitive neurons following 100% reward CSs. Before averaging, single neurons’ CS responses were normalized to the maximum CS response across 12 CS conditions; from 0 to 1. Error bars represent SEM. Ns – not significant (paired sign rank test; p>0.05).

**Supplemental Figure 12.**
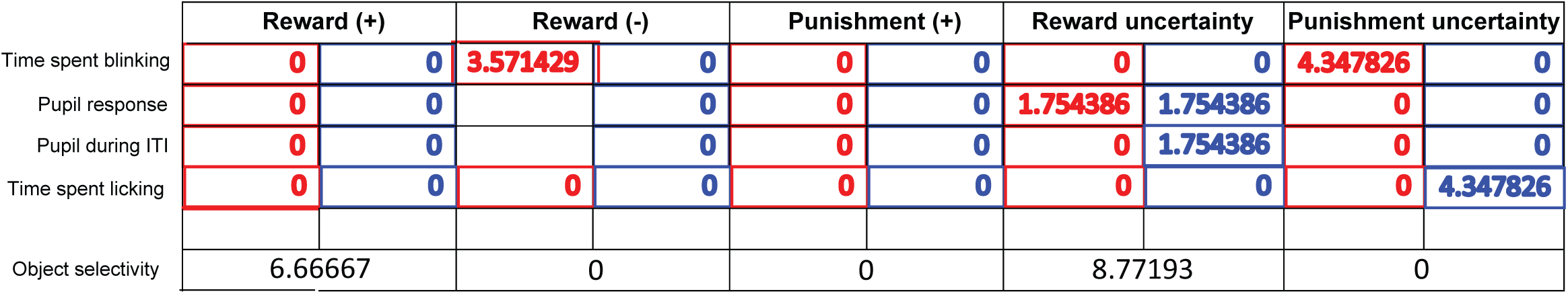
Summary of trial-by-trial relationships of ACC neurons with conditioned responses. Neurons were categorized into types (top of table) based on the results of Figures 2 and 4. The percentage of neurons in each category that displayed trial-by-trial correlations with conditioned responses (left) is reported. Positive correlations are colored red, negative are colored in blue. Below, percentage of neurons that displayed a difference in their responses to two different visual fractal objects that conveyed the same outcome probability is reported. Significance thresholds are p<0.01.

## Online Methods

### General procedures

Three adult male rhesus monkeys (*Macaca mulatta*) were used for the neurophysiology experiments (Monkeys B and Z and R). All procedures conformed to the Guide for the Care and Use of Laboratory Animals and were approved by the Washington University Institutional Animal Care and Use Committee. A plastic head holder and plastic recording chamber were fixed to the skull under general anesthesia and sterile surgical conditions. The chambers were tilted laterally by 35° and aimed at the anterior cingulate. After the monkeys recovered from surgery, they participated in the behavioral and neurophysiological experiments.

### Data acquisition

While the monkeys participated in the behavioral procedure we recorded single neurons in the anterior cingulate. The recording sites were determined with 1 mm-spacing grid system and with the aid of MR images (3T) obtained along the direction of the recording chamber. This MRI-based estimation of neuron recording locations was aided by custom-built software (PyElectrode). Single-unit recording was performed using glass coated electrodes (Alpha Omega). The electrode was inserted into the brain through a stainless-steel guide tube and advanced by an oil-driven micromanipulator (MO-97A, Narishige). Signal acquisition (including amplification and filtering) was performed using Alpha Omega 44 kHz SNR system. Action potential waveforms were identified online by multiple time-amplitude windows with an additional template matching algorithm (Alpha-Omega). Neuronal recording was restricted to single neurons that were isolated online. Neuronal and behavioral analyses were conducted offline in Matlab (Mathworks, Natick, MA).

Eye position was obtained with an infrared video camera (Eyelink, SR Research). Behavioral events and visual stimuli were controlled by Matlab (Mathworks, Natick, MA) with Psychophysics Toolbox extensions. Juice, used as reward, was delivered with a solenoid delivery reward system (CRIST Instruments). Juice-related licking was measured and quantified using previously described methods (62). Airpuff (∼35 psi) was delivered through a narrow tube placed 6–8 cm from the monkey’s face.

### Behavioral procedures

The reward-punishment behavioral procedure consisted of two alternating trial blocks: reward block and punishment block. In the reward block, three visual fractal conditioned stimuli (CS) were followed by a liquid reward (0.4 ml of juice) with 100%, 50%, and 0% chance, respectively. In the punishment block, three visual fractal CSs were followed by an air puff with 100%, 50%, and 0% chance, respectively. One block consisted of 12 trials with fixed proportions of trial types (each of the three CSs appears 4 times during each block). The inter-trial-intervals (ITIs) ranged from ∼2-6s.

Each trial started with the presentation of a trial-start cue at the center. The trial-start cue disappeared after 1 s and one of the three CSs was presented pseudo randomly (the CS could appear in three locations: 10 degrees to the left or to the right of the trial-start cue, or in the center). After 2.5 s, the CS disappeared, and the outcome (juice, aifpuff, or nothing) was delivered. The monkeys were not required to fixate.

Abort cues (Figure 1D) were presented during 50% one half of the trials in distinct blocks (abort-blocks) in one of three locations 10 degrees away from the CS. Abort blocks contained distinct visual fractals as CSs. Typically, a recording session contained the following repeating block structure: reward (abort) block, punishment (abort) block, reward block, punishment block, reward block, punishment block. Monkeys R, B, and Z participated in this procedure.

An additional control single block reward-punishment behavioral procedure was used to study the activity of ACC in monkey 1. In this study, the trial structure was the same as in the two block condition. Here, 9 visual fractals served as CSs that predicted reward with 100, 75, 50, 25, 0% chance and punishment with 100, 75, 50, and 25% chance. No abort cues were presented in this procedure. Monkey B participated in this procedure.

To study if reward-uncertainty sensitive ACC neurons signal the level of reward uncertainty and reward size, a five reward-probability and reward-amount procedure was used. This procedure consisted of two blocks, a reward-probability block and a reward-amount block. The trial structure was the same as in other experiments in this study and was detailed in White and Monosov, 2017. Each trial started with the presentation of a green trial-start cue at the center. The monkeys had to maintain fixation on the trial-start cue for 1 s; then the trial start cue disappeared and one of the CSs was presented pseudo randomly. After 2.5 s, the CS disappeared, and juice (if scheduled for that trial) was delivered. The reward-probability block contained five objects associated with five probabilistic reward predictions (0, 25, 50, 75 and 100% of 0.25 ml of juice) and a reward-amount block that contained five objects associated with 100% reward predictions of varying reward amounts (0.25, 0.1875, 0.125, 0.065 and 0ml). One block consisted of 20 trials with fixed proportions of trial types (each of the five CSs appears four times each block). Monkeys R, B, and Z participated in this procedure.

In separate experimental sessions, the monkeys’ choice preference was tested for the CSs. This procedure has been detailed in our previous studies (47, 62). Each trial started with the presentation of the trial-start cue at the center, and the monkeys had to fixate it for 1 s. Then two CSs appeared 10 degrees to the left and right. The fixation spot disappeared 500 ms after the appearance of the CSs, and the monkey had to make a saccade to one of the two CSs within 5 s and fixate it for at least 750 ms. Then the unchosen CS disappeared, and after 1s the outcome (associated with the chosen CS) was delivered and the chosen CS disappeared. If the monkey failed to fixate one of the CSs, the trial was aborted and all stimuli disappeared. The trials were presented pseudo randomly, so that a block of 180 trials contained all possible combinations of the 10 CSs four times. Monkeys R, B, and Z participated in this procedure.

### Data processing and statistics

Spike-density functions (SDFs) were generated by convolving spike-times with a Gaussian filter (σ = 100ms). All statistical tests were two-tailed. For comparisons between two task conditions for each neuron, we used a rank-sum test, unless otherwise noted. For comparisons between two task conditions across the population average we used a paired signed-rank test, unless otherwise noted. All correlation analyses were Spearman’s rank correlations. The significance of the correlation analyses (p<0.05) was tested by 10,000 permutations (41, 46, 47, 62).

CS responses were measured in a single time window that started 100 ms from the CS onset until the outcome. Outcome responses were measured in a single 500 ms window that started 50 ms following outcome delivery. To normalize task-event related responses, we subtracted baseline activity (the last 500 milliseconds of the inter-trial interval) from the activity during the task-event related measurement epoch.

A neuron was defined as CS responsive if it displayed variance across CSs either in the reward or punishment block (Kruskal-Wallis test, *P* < 0.01). The statistical identification of uncertainty neurons was detailed previously (41, 46, 47, 62). As before, a neuron was defined as uncertainty sensitive if its CS responses varied across the 3 possible outcome predictions in either block (Kruskal-Wallis test, *P* < 0.01) and if its response to the uncertain CS (50%) was significantly stronger or weaker than its responses to both 100% and 0% CSs (two-tailed rank-sum test; *P* < 0.05; Bonferroni corrected). Therefore, a neuron could be uncertainty sensitive in reward block, punishment block, or both (Figure 3B).

To calculate receiver operating characteristic (ROC) that assessed neuronal discrimination of value and uncertainty, we compared spike counts during the CS epoch of two conditions. The analysis was structured so that area under curve (AUC) values > 0.5 indicate that the response for the value or uncertainty was a selective enhancement, while values < 0.5 indicate that the response for the value or uncertainty was a selective suppression.

Trial-by-trial correlations of single unit activity and conditioned motor responses (Supplemental Figure 12) were performed so as to minimize the influence of global task correlations. For each neuron type, the neurons’ preferred CS was chosen (e.g. reward uncertainty enhanced neurons’ 50% reward CS epochs were analyzed), and the relationship of spiking activity (mean spike count) and the magnitude of conditioned responses during that CS epoch was tested by a Spearman’s rank correlation. Similar results were obtained if partial correlations were performed across all CS conditions in which spiking activity and conditioned responses were z-scored within each CS condition before the correlation was performed. For the trial-by-trial correlations assessing firing rate during the CS epoch and the subsequent inter-trial-interval pupil dilation, the pupils were assessed in a 0.50 s window, 1.5 s following the outcome.

For each neuron, object selectivity was assessed by comparing its responses to two distinct visual fractals that conveyed the neuron’s preferred outcome prediction (e.g. reward value enhanced neurons’ responses for two visual fractals that conveyed 100% reward).

The goal of Supplemental Figure 5 was to visualize how uncertainty and value coding strategies co-occur in single neurons across the reward and punishment blocks. Each neuron was represented by a vector of 4 value and uncertainty discrimination indices ranging from 0 to 1. Value discrimination indices were obtained by ROC analyses that compared 100% and 0% CS responses. Uncertainty discrimination indices were obtained by ROC analyses that compared 50% and 100% CS responses. If a neuron did not pass the uncertainty sensitivity test (described above), the result of the ROC analysis was set to 0.5. If a neuron was uncertainty sensitive in one of the blocks, its value index was set to 0.5 in that block. Hierarchical clustering organized these data into 8 clusters. This was done by clustering on Euclidean distance utilizing Ward’s minimum variance method. The number of clusters was approximated based on the results of expectation maximization algorithm (81). However, because clustering is a data mining method, here it is utilized to organize the matrix of neurons for qualitative observation. The results are interpreted as visual confirmation for multiple value and uncertainty coding strategies in ACC, not as evidence for discrete and stable clusters (e.g. robust to noise or increases or decreases in neuron number).

## Acknowledgements

This work was supported by the Defense Advanced Research Projects Agency (DARPA) Biological Technologies Office (BTO) ElectRx program under the auspices of Dr. Doug Weber through the CMO Grant/Contract No.HR0011-16-2-0022. I am grateful to Mr. J. Kael White, Ms. Kim Kocher, and Ms. Ying Jiang for assisting in data acquisition, to Ms. Kim Kocher for fantastic animal care and training, and to Ms. Katherine Conen, Mr. J. Kael White, Mr. Ben Acland, and Mr. Mike Traner for reading earlier versions of this manuscript.

